# Histone demethylase UTX regulates glioblastoma progression through affecting periostin expression

**DOI:** 10.1101/2021.12.17.473115

**Authors:** Yan Luan, Yingfei Liu, Jingwen Xue, Ke Wang, Kaige Ma, Haixia Lu, Xinlin Chen, Yong Liu, Zhichao Zhang

**Affiliations:** Institute of Neurobiology, Xi’an Jiaotong University Health Science Center, Xi’an, China; Department of Critical Care Medicine, the First Affiliated Hospital of Xi’an Jiaotong University, Xi’an, China; Department of Urology, the First Affiliated Hospital of Xi’an Jiaotong University, Xi’an, China

**Author notes:** These authors contributed equally to this work (Yan Luan, Yingfei Liu). **Correspondence:** Dr. Zhichao Zhang, Institute of Neurobiology, Xi’an Jiaotong University Health Science Center, 76 Yanta West Road, Xi’an, Shaanxi 710061, China.

**Keywords:** UTX, Glioblastoma, Tumor progression, Periostin, Patient-derived glioblastoma stem-like cells

## Abstract

The histone H3K27 demethylase UTX participates in regulating multiple cancer types. However, less is known about the UTX function in glioblastoma (GBM). This study aims to define the effect of UTX on GBM. GEPIA2 database analysis showed that UTX expression was significantly increased in GBM and inversely correlated with survival. Knockdown UTX inhibited GBM cell proliferation, migration, and invasion while promoting apoptosis. Moreover, knockdown UTX also hampered tumor growth in the heterotopic xenograft model. RNA-seq combined with qRT-PCR and ChIP-qPCR were used to identify the target genes. The results showed that the UTX-mediated genes were strongly associated with tumor progression and the extracellular environment. Protein-protein interaction analysis suggested that periostin (POSTN) interacted with most of the other UTX-mediated genes. POSTN supplement abolished the effect of UTX knockdown in GBM cells. Furthermore, silencing UTX exhibited similar antitumor effect in patient-derived glioblastoma stem-like cells, while UTX functions were partially restored after exposing POSTN. Our findings may reveal a new insight into the onset of gliomagenesis and progression, providing a promising therapeutic strategy for GBM treatment.

**Bullet Points:** UTX correlates with survival in glioblastoma. Silencing UTX decreased the levels of H3K27 methylation in the POSTN gene, thereby suppressing POSTN expression. UTX-mediated POSTN expression is crucial for glioblastoma cell growth and tumorigenesis.

1. UTX expression is increased in GBM and negatively correlated with survival.
2. UTX knockdown influences proliferation and apoptosis in both GBM cells and GSCs.
3. The antitumor effect of UTX knockdown is achieved by suppressing POSTN expression.

## Introduction

Glioblastoma (GBM), a WHO grade IV glioma, is the most common primitive malignant brain tumor, characterized by rapid progression, high metastasis and fast relapse (Davis, 2016). Generally, the treatment for GBM is surgery followed by chemotherapy, radiotherapy or both. Despite the efforts made in recent years, 5-year survival rates for patients with GBM are poor (nearly 3-5%) and the median survial is less than 15 months (Shergalis *et al*, 2018). In addition, glioblastoma stem-like cells (GSCs) redoubles the difficulty of treatment due to the greater resistance to currently antineoplastic agents (Dirkse *et al*, 2019). A cure for GBM therefore remains elusive. Continuing efforts are in need to explore and develop a novel effective therapeutic target for both GBM and GSCs.

The *Ubiquitously Transcribed Tetratricopeptide Repeat on chromosome X* (UTX, also called KDM6A) is a histone demethylase that catalyzes the removal of di- and tri-methyl marks at histone H3 lysine 27 (H3K27) (Agger *et al*, 2007). Previous studies indicated that UTX underpinned a substantial component of epigenetic deregulation and it was strongly associated with multiple types of human cancer (Kobatake *et al*, 2020; Leng *et al*, 2020; van Haaften *et al*, 2009). UTX inhibited the proliferation, invasion, and metastasis of breast cancer cells by interacting with GATA3 (Yu *et al*, 2019). For lung cancer, UTX deletion dramatically promoted tumorigenesis and progression (Wu *et al*, 2018). In contrast, H3K27 methyltransferase EZH2 has a tumor-promotive effect. It is highly expressed in several forms of cancer and regulates various oncogenic transcription factors, tumor suppressor miRNAs, and cancer-associated non-coding RNA (Yamaguchi & Hung, 2014). Although the exact role of UTX in GBM is poorly understood at present, inhibition of JMJD3 (also called KDM6B, an H3K27me3 demethylase) suppressed glioma cell proliferation, migration, and promoted cell apoptosis (Sui *et al*, 2017). These phenomena suggest UTX might involve in the regulation of GBM progression.

The tumor microenvironment (TME) is comprised by tumor cells, blood vessels, other non-malignant cells, and extra-cellular components (Wu & Dai, 2017). Accumulating evidence suggests that TME is a novel target in tumor therapy (Allegrezza & Conejo-Garcia, 2017). Periostin (POSTN, also known as OSF-2), one of the components of extracellular matrix, belongs to the fasciclin family and is involved in regulating tissue/organ development, pathological fibrosis, tissue remodeling and cancer biology progression (Izuhara et al, 2014; Li et al, 2015). POSTN is secreted by different cells types in solid tumors and exerts its functions through autocrine and paracrine. Secreted POSTN can activate invasion- and survival-related signaling pathways, which in turn promotes invasion, proliferation and survival (Ruan *et al*, 2009). For GBM, POSTN expression level was correlated with tumor grade and recurrence. Overexpression of POSTN promoted glioma cell invasion and apoptosis (Mikheev *et al*, 2015). More importantly, POSTN was among the most upregulated genes in GBM compared to the normal brain tissue (Tso *et al*, 2006). Silencing POSTN in glioma stem cells inhibited cell proliferation and increased survival of xenografts mice (Zhou *et al*, 2015a). These results suggested that POSTN might be an excellent target for GBM therapeutics. However, the mechanisms of regulation of POSTN expression require further investigation.

In this study, we analyzed the relationship between the expression level of UTX and GBM malignancy. The effects and underlying molecular mechanisms of UTX in both GBM cells and patient-derived GSCs were further investigated. Here we found that high UTX expression promoted both GBM progression and GSCs proliferation. Moreover, our present study may provide evidence that the suppression UTX decreased the methylation of H3K27 in the POSTN gene, thereby suppressing POSTN expression which ultimately inhibits growth and tumorigenesis. These results may reveal new insight into the onset of gliomagenesis and progression, provide a vigorous therapeutic strategy for GBM treatment.

## Results

### UTX knockdown restricts GBM cell proliferation

The GEPIA2 databases (including normal, low-grade glioma (LGG) and glioblastoma multiform samples) was used to analyze the differential expression of UTX. Compared to the normal group, the UTX expression level was significantly increased in GBM and oligodendrogliomas (ODs, a subcategory of LGG) (Fig. 1 A, B). Moreover, UTX expression was negatively correlated with overall survival for LGG and GBM patients (Fig. 1 C). These results suggested that UTX was a risk factor and blockade of UTX expression might be developed for GBM treatment.

**Figure 1.**
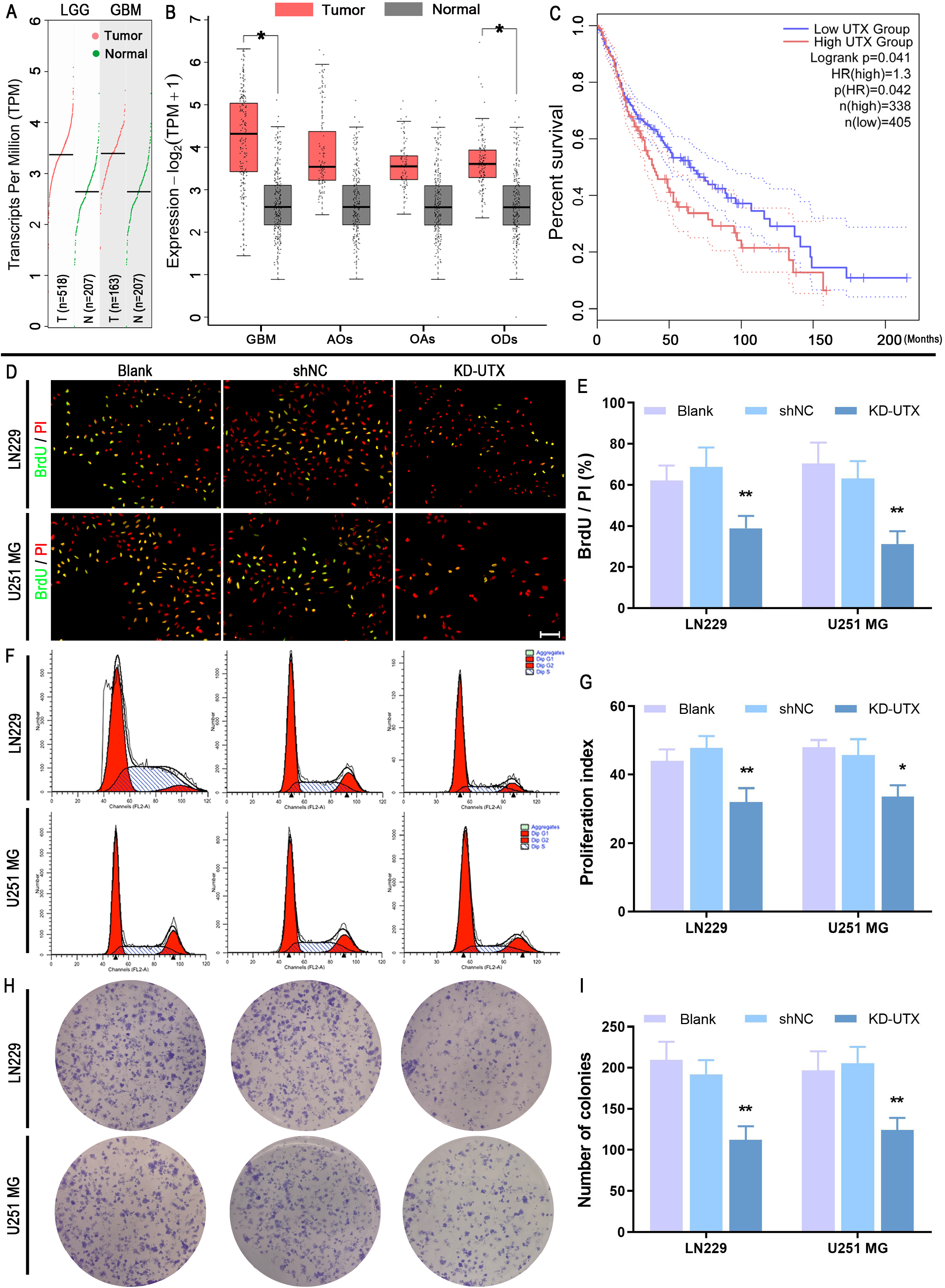
UTX knockdown inhibits the ability of proliferation in GBM cells. The clinical information data for normal (n=207), LGG (n=518) and GBM (n=163) were obtained and analyzed by GEPIA 2 (http://gepia2.cancer-pku.cn/#index). (A, B) The expression level of UTX in normal tissue, LGG and GBM were present as transcripts per million or log2 transcripts per million. (C) Kaplan-Meier survival analysis was performed for overall survival of LGG and GBM patients. The infected LN229 and U251 MG cells were divided into three groups. Blank group: cells were maintained without any treatment; shNC group: cells infected with nonspecific shRNA lentivirus; KD-UTX group: cells infected with anti-UTX shRNA lentivirus. (D) After culturing 3 days, BrdU-positive cells were determined by immunostaining, and the result was shown as percentages among PI-stained cells. Scale bar 100 μm (F) Proliferation index was determined by cell cycle analysis. (H) The colony formation assay was performed to investigate the long-term effect of UTX on proliferation. (E, G and I) Data are presented as the mean ± standard deviation of three independent experiments (n = 3). \**p* < 0.05, \*\**p* < 0.01 versus the shNC group. LGG, Low-grade glioma; GBM, glioblastoma; ODs, oligodendrogliomas; AOs, astrocytomas; OAs, oligoastrocytomas.

To specifically inhibit UTX expression, LN-229 and U251 MG cell lines were transfected with three UTX-target siRNAs (si-UTX-1/2/3). The combined results from RT-PCR and Western blot indicated that si-UTX-1 exerted a higher inhibition efficiency (Fig. EV1 A-E). Therefore, the si-UTX-1 sequence was used in the subsequent lentiviral packaging. The lentivirus-transduced cells were selected using puromycin. We found that the vast majority of cells expressed eGFP and the expression of UTX was significantly decreased (Fig. EV1 F-J). This lentiviral UTX shRNA (KD-UTX) was used for further experiments. The proliferation effect of UTX on GBM was verified by BrdU incorporation and cell cycle analysis. The results showed that there was no significant difference between the blank group (cells without any treatment) and the shNC group (cells infected with nonspecific control shRNA). Compared to the shNC group, UTX knockdown significantly decreased the number of BrdU-positive cells and proliferation index (Fig. 1 D-G). To further verify the role of UTX in GBM cell proliferation, we performed a colony formation assay to investigate the long-term effect of UTX on proliferation. As we except, the colony-forming ability was decreased in both LN-229 and U251 MG cell lines after UTX knockdown treatment (Fig. 1 H, I). These data suggest that UTX knockdown impairs the proliferation of GBM cells.

### UTX knockdown affects GBM cell apoptosis, migration and invasion

TUNEL staining and flow cytometry with Annexin V/PI were applied to distinguish apoptotic cells. There was an increase in the number of TUNEL-positive cells following UTX knockdown treatment (Fig. 2 A-C). Similar to the results of TUNEL assay, the apoptosis rate was significantly increased in the KD-UTX group (Fig. 2 D-F). Then, wound healing and transwell invasion assay were used to detect the migration and invasion abilities in GBM cells, respectively. Wound healing assay showed that UTX knockdown markedly attenuated the migration capacity of LN229 and U251 MG cells (Fig. 2 G-I). The result of the transwell invasion assay showed that the number of transwell invasive cells was significantly decreased by suppressing the UTX expression in GBM cells (Fig. 2 J-L). Taken together, these data indicate that UTX may promote migration and invasion while inhibiting apoptosis in GBM cells.

**Figure 2.**
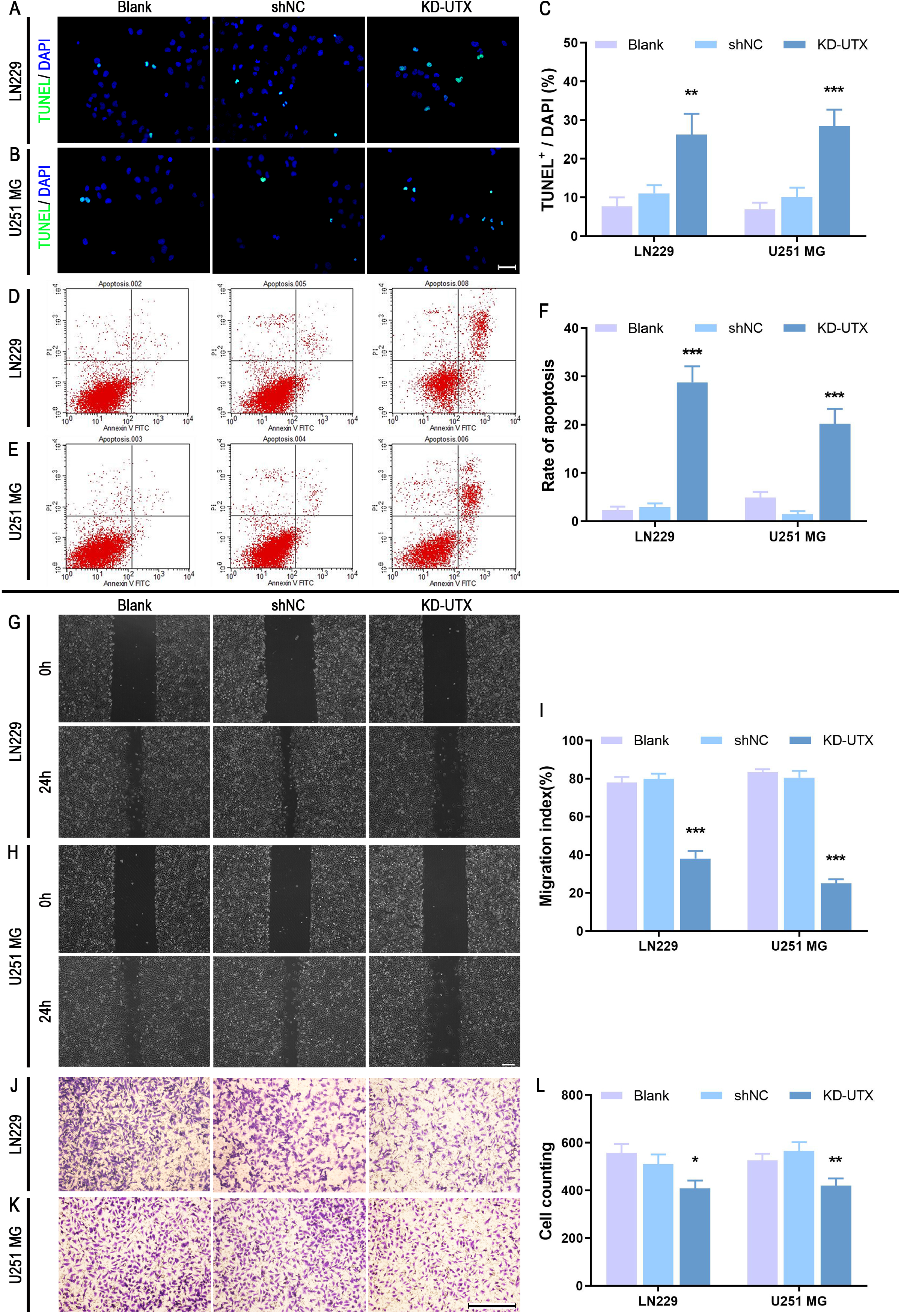
UTX knockdown promotes GBM cell apoptosis, inhibits migration and invasion. The infected LN229 and U251 MG cells were divided into three groups. Blank group: cells were maintained without any treatment; shNC group: cells infected with nonspecific shRNA; KD-UTX group: cells infected with anti-UTX shRNA. After culturing 3 days, TUNEL staining (A, B) and flow cytometry analysis (D, E) were performed to evaluate cell apoptosis. (G, H) Representative images of scratch-wound healing exhibit the motility of GBM cells. (J, K) Cell invasion was assessed by Transwell chambers coated with Matrigel. (C, F, I and L) Data are presented as the mean ± standard deviation of three independent experiments (n = 3). \**p* < 0.05, \*\**p* < 0.01, \*\*\**p* < 0.001 versus the shNC group. Scale bar, 50 μm (A, B), 500 μm (G, H); 200 μm (J, K).

### UTX knockdown inhibits tumor growth in a heterotopic glioblastoma xenograft model

The stable UTX knockdown LN229 cells were implanted into the subcutis of the athymic nude mice to establish the heterotopic xenograft model. We measured the tumor volumes and weights, and detected the expression of apoptosis related Caspase-3 and Cyclin D1 at different time points. Tumor volumes and weights were significantly decreased in the LN229-KD-UTX group (Fig. 3 A-C). Furthermore, we found Cleaved-caspase-3 / Pro-caspase-3 ratio was increased, while the expression of CyclinD1 was decreased (Fig. 3 D-F). These results obtained with the heterotopic glioblastoma xenograft model were consistent with the *in vitro* data, suggesting that UTX may be involved in the regulation of GBM cell proliferation and apoptosis.

**Figure 3.**
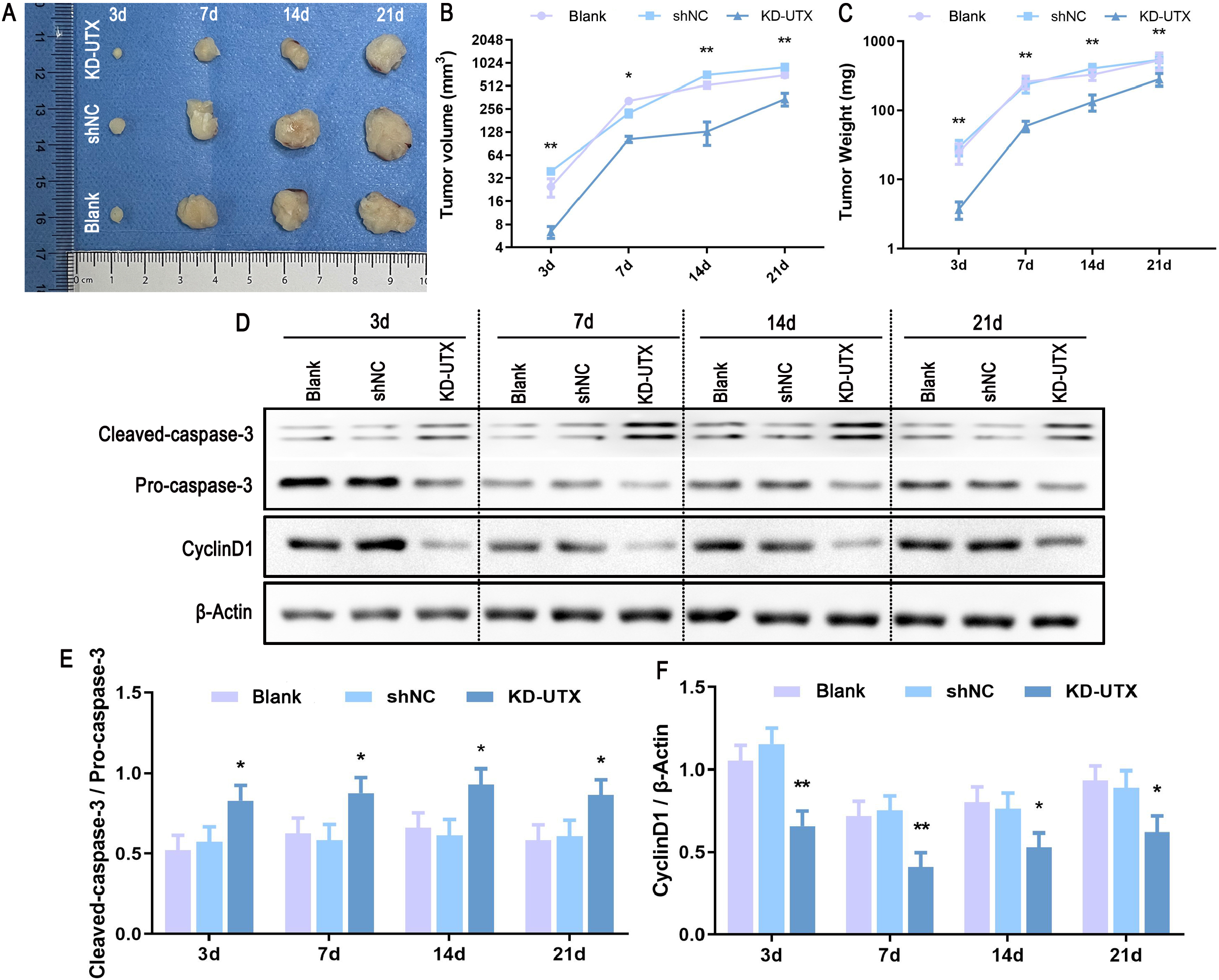
UTX knockdown influences the proliferation and apoptosis of LN229 cells in a heterotopic xenograft mouse model. The stable UTX knockdown LN229 cells (1 × 10^6^ per injection) were implanted subcutaneously into the nude mice in the heterotopic xenograft experiment. As negative control, shNC LN229 cells (1 × 10^6^ per injection) were implanted subcutaneously into the nude mice. For Blank group, 1 × 10^6^ of normal LN229 cells were implanted subcutaneously into the nude mice. Mice were killed at 3, 7, 14, and 28 days (the third day after cell inoculation was deemed as 0 d). Five mice of each group were killed at each timepoint, tumors were removed for (A) photograph, the measure of the tumor (B) volumes (length×width^2^×0.52) and (C) weights, and (D) Western blotting. (E) Immunoblot quantification for the ratio of cleaved-caspase-3 to pro-caspase-3. (F) Immunoblot quantification for the ratio of cyclin D1 to β-Actin. Data are presented as the mean ± standard deviation of five independent experiments (n = 5). \**p* < 0.05, \*\**p* < 0.01 versus the shNC group.

### UTX knockdown influences the expression of various secreted protein-coding genes

To gain insight into the consequences of UTX knockdown in the LN229 cells, the whole transcriptome analysis was performed by using RNA-seq after culturing for 3 days. Principal component analyses (PCA) showed that the transcriptome of UTX knockdown cells was clearly segregated from the shNC group (Fig. EV2 A). 469 genes expression were significantly altered by UTX withdrawal. Of these, 374 genes expression increased, while 95 genes decreased (Fig. 4 A, Fig EV2 B, C). Geneset enrichment analysis (GSEA) showed DNA replication, cell cycle checkpoint, G1/S and G2/M phase transition genes were significantly enriched in the downregulated genes (Fig. 4 B). Gene Ontology (GO) enrichment analysis showed that most of the altered genes were preferentially correlated with secretion, extracellular matrix and extracellular structure organization (Fig. 4 C). Moreover, we underwent Disease Ontology (DO) (Fig. 4 D) and KEGG analysis (Fig. EV2 D), and the data showed that the changes in gene expression were closely associated with cancers. These phenomena suggested that UTX may regulate the onset and progression of tumors through influencing the extracellular matrix.

**Figure 4.**
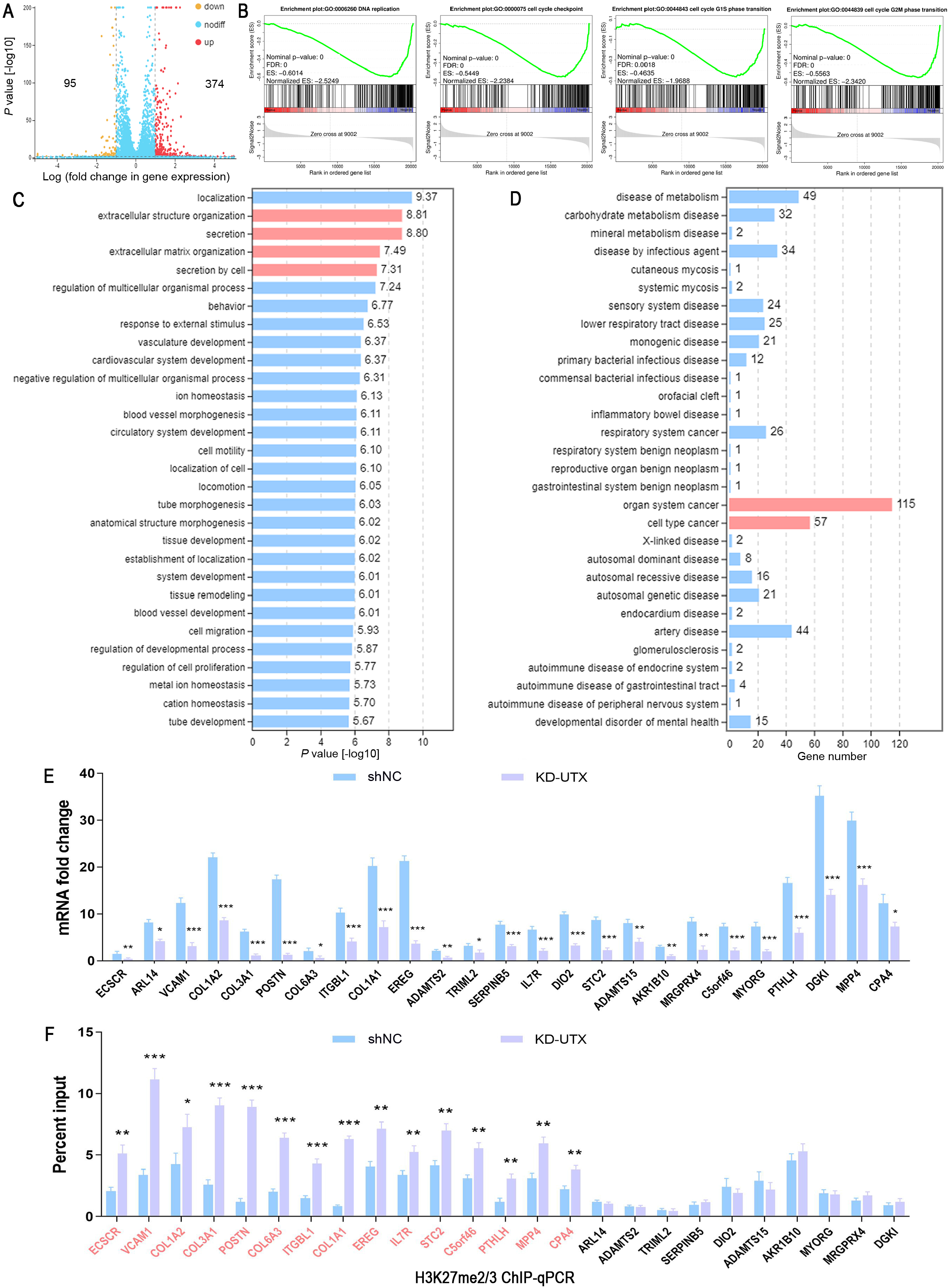
RNA-Seq reveals alteration of gene expression after UTX knockdown in LN229 cell line. LN229 cells with stable expression of UTX-shRNA (KD-UTX) or scramble shRNA (shNC) were cultured for 3 days before total RNA collection. (A) Volcano plots showed global differences between shNC and KD-UTX cells. Gene-set enrichment analysis (GSEA) (B), Gene Ontology (GO) enrichment (C) and Disease Ontology (DO) analysis (D) were performed to interpret the results. (E) To experimental validation of RNA-seq data, significantly downregulated genes (>1.5 fold) were selected and detected the expression by qRT-PCR. (F) ChIP-qPCR was used to measure the H3K27me2/3 enrichment in the significantly downregulated genes. The genes with a significantly elevated level of H3K27me2/3 were labeled in red. Data are presented as the mean ± standard deviation of three independent experiments (n = 3). \**p* < 0.05, \*\**p* < 0.01, \*\*\**p* < 0.001 versus the shNC group.

UTX is linked with the demethylation of lysine residues on H3K27, so we detected the level of H3K27me2/3 by Western blot. As expected, the level of H3K27me2/3 was significantly increased in the KD-UTX group (Fig. EV2 E, F). H3K27me2/3 is usually associated with gene repression, thus we focused on the 95 downregulated genes. The significantly (>1.5 fold) downregulated genes were selected and qRT-PCR was carried out to validate the sequencing results. Most qRT-PCR results of the genes were consistent with that of RNA-seq (Fig. 4 E), while the expression of some genes was not significantly decreased or slightly increased (Fig. EV2 G). ChIP-qPCR was carried out to further confirm the regulation of H3K27me2/3 on these downregulated genes. The results revealed that the level of H3K27me2/3 was increased in 15 genes (Fig. 4 F). Interestingly, most of the genes, including POSTN. VCAM1 and COL1A1, are primarily responsible for encoding secreted proteins that in turn modulate the extracellular microenvironment.

### UTX knockdown affects many extracellular matrix proteins expression

To further investigate the effect of UTX on the expression of extracellular matrix protein, intracellular protein and medium-containing protein were detected by Western blot and ELISA, respectively. POSTN, COL1A1 and VCAM1 concentrations in the culture medium were significantly decreased by UTX knockdown treatment (Fig. 5 A, B). Moreover, Western blot showed that the levels of intracellular proteins (including POSTN, COL1A1, PTHLH, VCAM1 and CPA4) were reduced as well (Fig. 5 C-E). The above results indicate that the effect of UTX on GBM cells may be mediated through regulating the tumor microenvironment.

**Figure 5.**
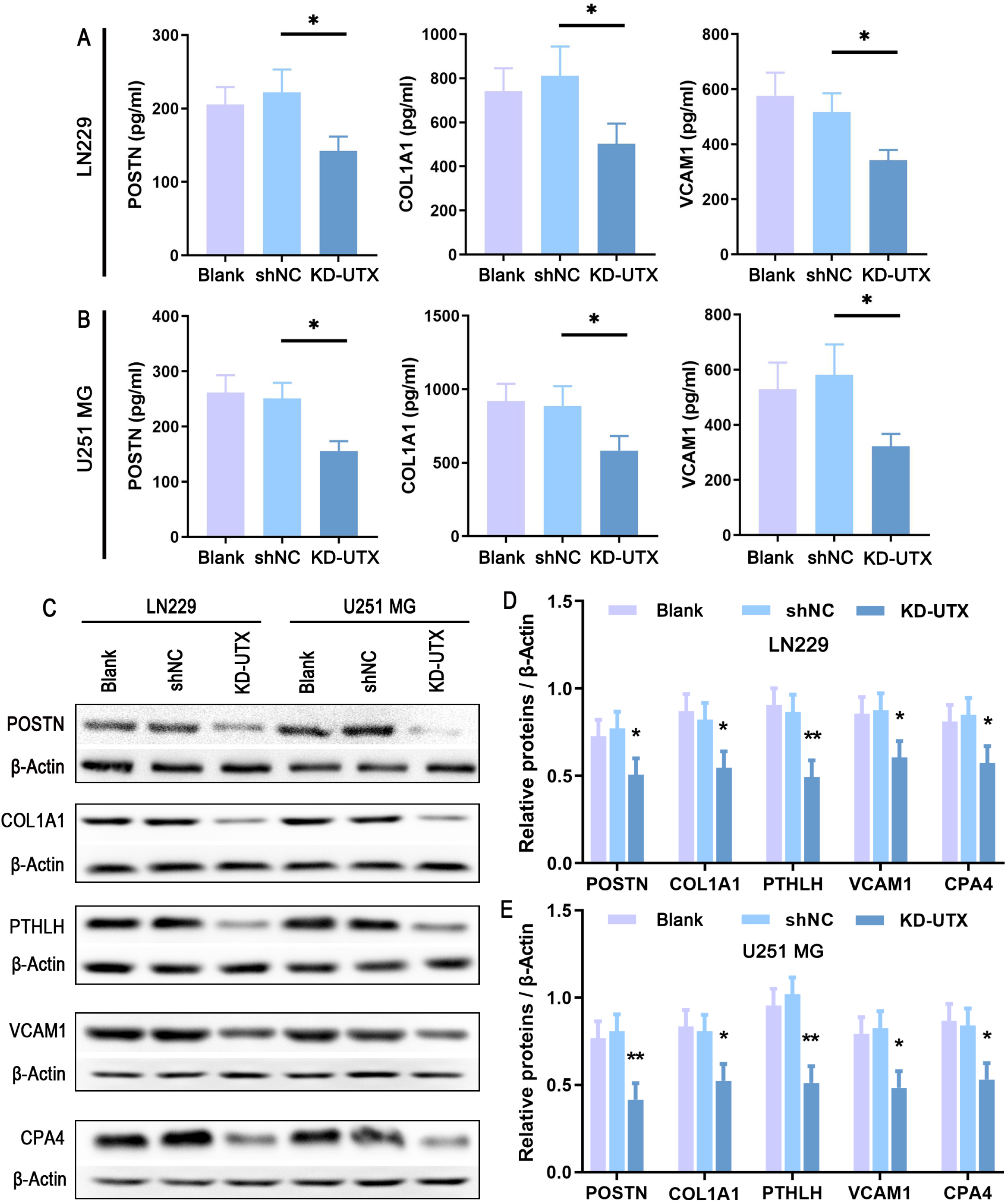
UTX knockdown inhibits the expression of extracellular matrix proteins. The infected LN229 and U251 MG cells were divided into three groups. Blank group: cells were maintained without any treatment; shNC group: cells infected with nonspecific shRNA; KD-UTX group: cells infected with anti-UTX shRNA. After culturing 3 days, cell lysates and culture medium were respectively collected and examined by ELISA and Western blot. (A, B) POSTN, COL1A1 and VCAM1 concentration in the culture medium were detected by ELISA. The values are presented as the mean ± standard deviation of three independent experiments (n = 3). \**p* < 0.05 versus shNC group. (C) The intracellular concentration of POSTN, COL1A1, PTHLH, VCAM1 and CPA4 were examined by Western blot. (D, E) Band intensity was quantified and plotted as a ratio of the target protein to β-Actin. The values are presented as the mean ± standard deviation of three independent experiments (n = 3). \**p* < 0.05, \*\**p* < 0.05 versus shNC group.

### POSTN is involved in the regulation of UTX on GBM progression

The proteins, which are regulated by H3K27me2/3, were analyzed by the STRING database to reveal protein-protein connections. We found that POSTN protein could interact with various proteins and might play a pivotal role in the transcriptional regulatory network (Fig. 6 A, B). To evaluate the involvement of POSTN in the regulation of UTX on GBM progression, 2 μg/mL recombinant human POSTN protein (R&D Systems, USA) was replenished in the KD-UTX cell culture medium. BrdU incorporation and CyclinD1 expression showed that the effect of UTX on cell proliferation was abolished after exposing POSTN (Fig. 6 C, D, G, H). Cellular apoptosis, detecting by TUNEL staining and caspase-3 activation, suggested that POSTN protein eliminated the pro-apoptotic effect of UTX knockdown in LN229 cells (Fig. 6 E, F, I, J). Furthermore, addition of POSTN protein partially restored the inhibitory effect of UTX knockdown on migration and invasion (Fig. 6 K-N). These data suggested that knockdown UTX promoted GBM progression by inhibiting POSTN expression

**Figure 6.**
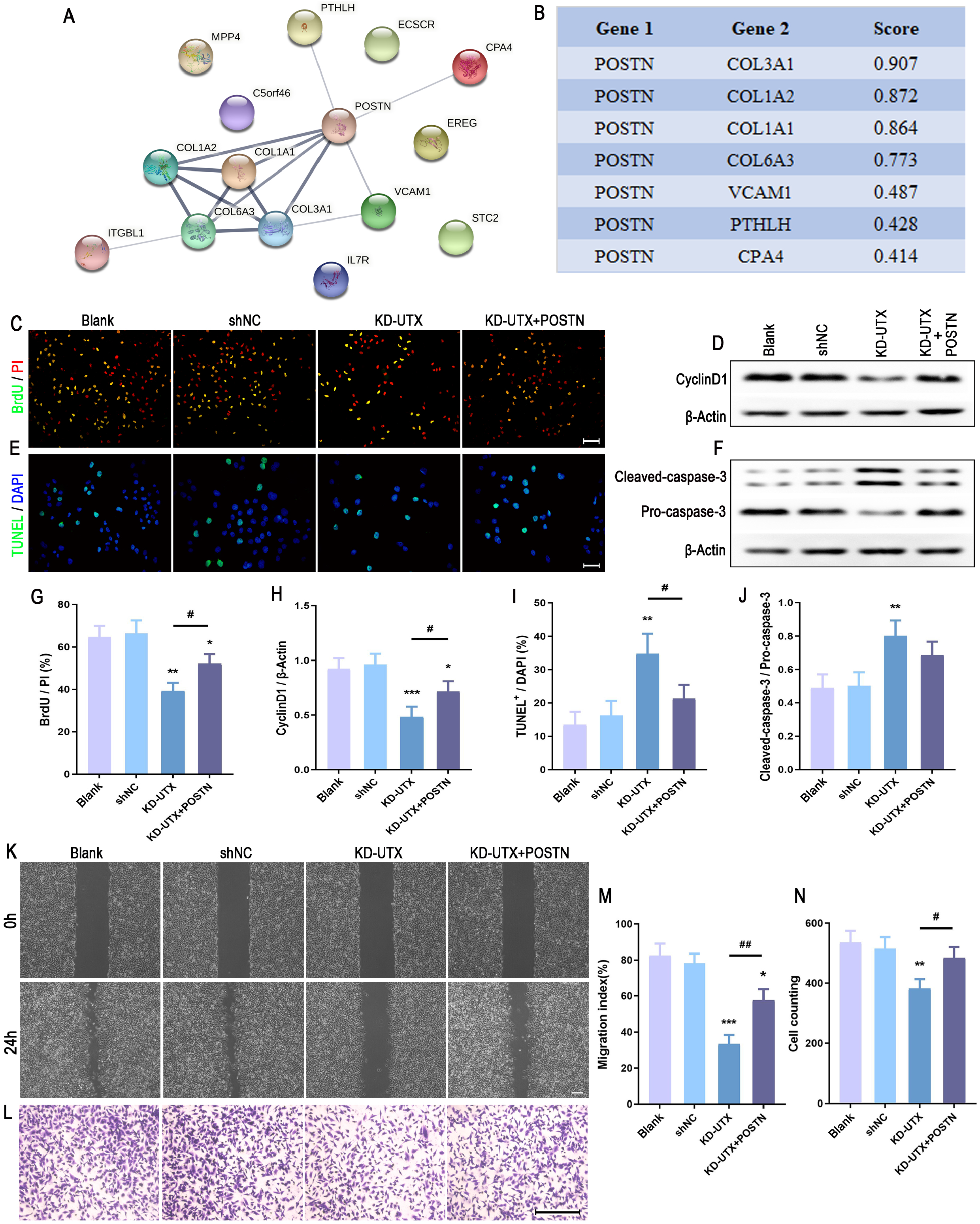
POSTN abolishes the effect of UTX on LN229 cell proliferation, apoptosis, migration and invasion. (A) Protein interactions (grey lines, the confidence score ≥ 0.4) were extracted from the STRING database. (B) The table showed the STRING score of POSTN interacts with other proteins. The infected LN229 cells were divided into four groups. Blank group: cells were maintained without any treatment; shNC group: cells infected with nonspecific shRNA; KD-UTX group: cells infected with anti-UTX shRNA; KD-UTX +POSTN group: added 2 μg/mL POSTN protein in the knockdown UTX cells culturing medium. BrdU incorporation was measured by immunostaining (C) and the expression of CyclinD1 was examined by Western blot (D). Cell apoptosis was detected by TUNEL assay (E) and the ratio of cleaved-caspase-3 to pro-caspase-3 (F). Wound healing (K) and transwell invasion assay (L) were evaluated the ability of migration and invasion, respectively. (G-J, M and N) The values are presented as the mean ± standard deviation of three independent experiments (n = 3). \**p* < 0.05, \*\**p* < 0.01, \*\*\**p* < 0.001 versus shNC group. #*p* < 0.05, ##*p* < 0.01 versus KD-UTX group. Scale bar, 100 μm (C); 50 μm (E); 500 μm (K); 200 μm (L).

### UTX knockdown influences proliferation and apoptosis of patient-derived GSCs by regulating POSTN expression

Previous studies demonstrated that POSTN played a crucial role in maintaining cancer stem cells and promoting tumor progression including growth, invasion and metastasis (Gillan *et al*, 2002; Michaylira *et al*, 2010). In the present study, we investigated whether UTX participates in regulating GSCs. At first, we isolated and cultured patient-derived GSCs. Human GSCs were isolated from three post-surgical patients (GSC02, GSC05 and GSC08). After culturing 5-7 days, 80-200 μM spheres were formed (Fig. 7 A). The double immunofluorescent staining showed that these spheres expressed cancer stem cell markers, including CD133, CD15, CD44 and nestin (Fig. 7 B, C). Moreover, the single-GSCs staining showed that 96.89% ± 5.37% nestin-positive cells, of which 96.65% ± 4.28% expressed SOX2 (Fig. 7 D). These phenomena suggested that these cultured cells were human GSCs, and these cells were used to conduct subsequent experiments. After infecting with KD-UTX lentivirus, the UTX expression was significantly decreased in the patient-derived GSCs (Fig. EV3 A, B). Moreover, silencing UTX inhibited POSTN protein (Fig. EV3 C, D) and mRNA expression (Fig. EV3 E) while enhancing the level of H3K27me2/3 (Fig. EV3 F). These results indicated that UTX regulated POSTN expression through altering H3K27 methylation levels in the patient-derived GSCs. To confirm the effect of UTX on GSCs, we measured cell viability by CCK-8 assay at different time-points. The results showed that low UTX expression in GSCs was characterized by a low proliferation rate (Fig. 7 E-G). Moreover, suppression of UTX resulted in a decreased number of BrdU-labelled cells (Fig. 7 H, G; Fig. EV4 A, C and E). In contrast, UTX knockdown significantly increased the TUNEL positive cells (Fig. 7 I, K; Fig. EV4 B, D and F). Interestingly, supplement POSTN in the medium abolished the effect of UTX on proliferation and apoptosis in GSCs. These data suggest that UTX can regulate the proliferation and apoptosis of the patient-derived GSCs by regulating POSTN expression.

**Figure 7.**
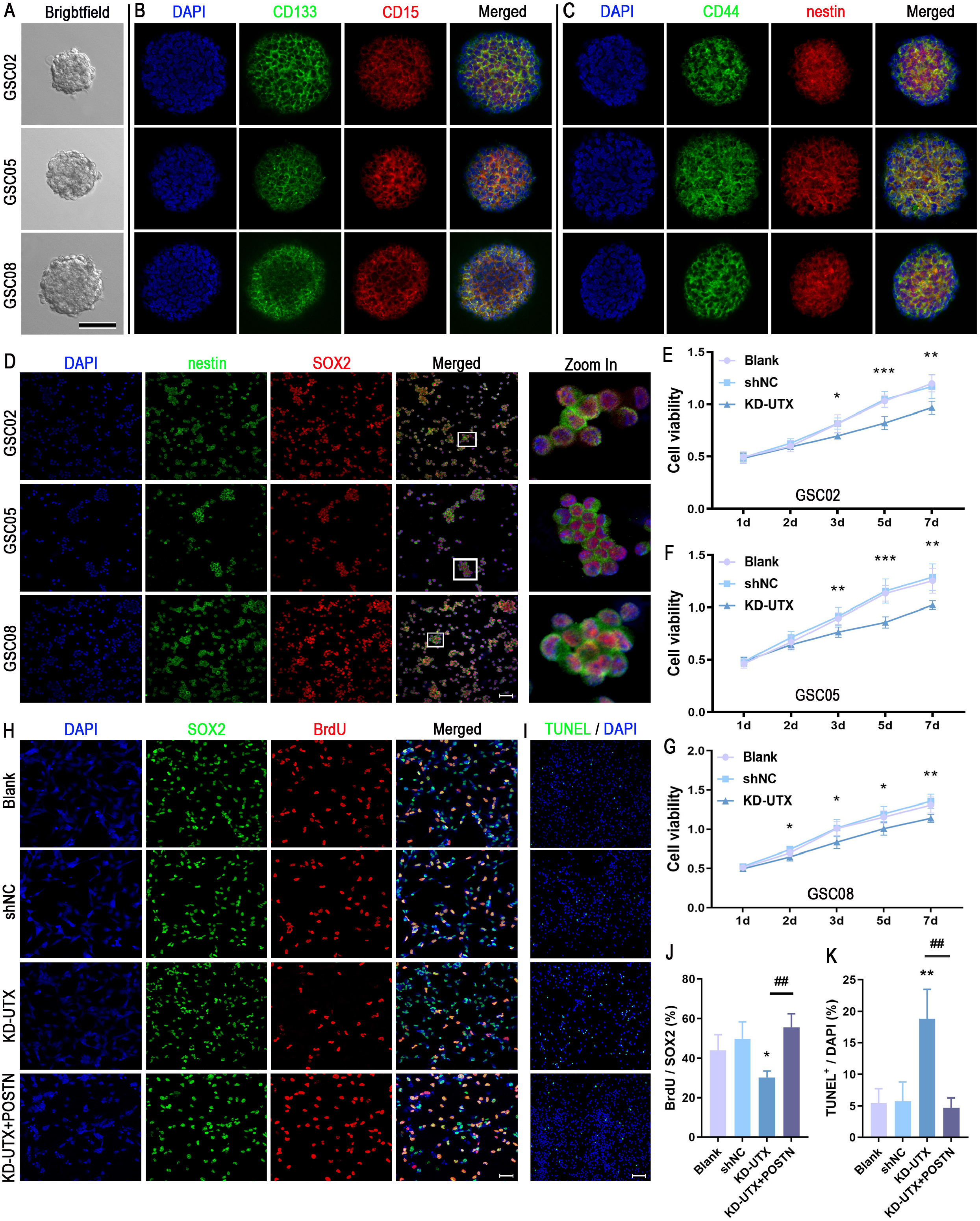
UTX knockdown displays an inhibiting role in patient-derived-GSCs through regulating POSTN. Human GSCs were isolated from three post-surgical patients (GSC02, GSC05 and GSC08). (A) Spheres, 80–120 μm in size were observed after culturing for 5-7 days. The spheres were identified by double immunofluorescent labeling for specific markers CD133 and CD15 (B) or CD44 and nestin (C). Patient-derived GSCs were infected with shNC or KD-UTX lentivirus and divided into four groups. Blank group: cells were maintained without any treatment; shNC group: cells infected with nonspecific shRNA; KD-UTX group: cells infected with anti-UTX shRNA; KD-UTX +POSTN group: added 2 μg/mL POSTN protein in the knockdown UTX cells culturing medium (E-G). The cell viability was quantitatively measured by using a CCK-8 at different time points (1-7 days). After culturing 3 days, cell proliferation was identified by BrdU staining (H) and apoptotic cells were detected by TUNEL staining (I). (J) Quantitative data from three independent experiments (n = 3) were shown as the percentage of BrdU-positive cells in total SOX2-stained cells. (K) Quantitative data from three independent experiments (n = 3) are presented as the percentage of TUNEL-positive cells in the total DAPI cells. \**p* < 0.05, \*\**p* < 0.01 versus shNC group; ##*P* < 0.01 versus KD-UTX group. Scale bar, 100 μm (A), 50 μm (D, H); 100 μm (I).

## Discussion

In this study, we showed that UTX expression was significantly elevated in GBM and was inversely correlated with survival. Knockdown UTX in both GBM cells and patient-derived GSCs decreased the methylation of H3K27 in the POSTN gene, thereby suppressing POSTN expression. The low expression level of POSTN affected lots of extracellular components (including COL1A1, VCAM1, PTHLH, and CPA4), which in turn inhibited growth and tumorigenesis. Moreover, supplement POSTN abolished the antitumor effect of UTX knockdown (Fig. 8). These results suggested that UTX was an oncogenic factor and could promote growth and tumorigenesis, and UTX inhibition was a novel therapeutic target for rational GBM drug development.

**Figure 8.**
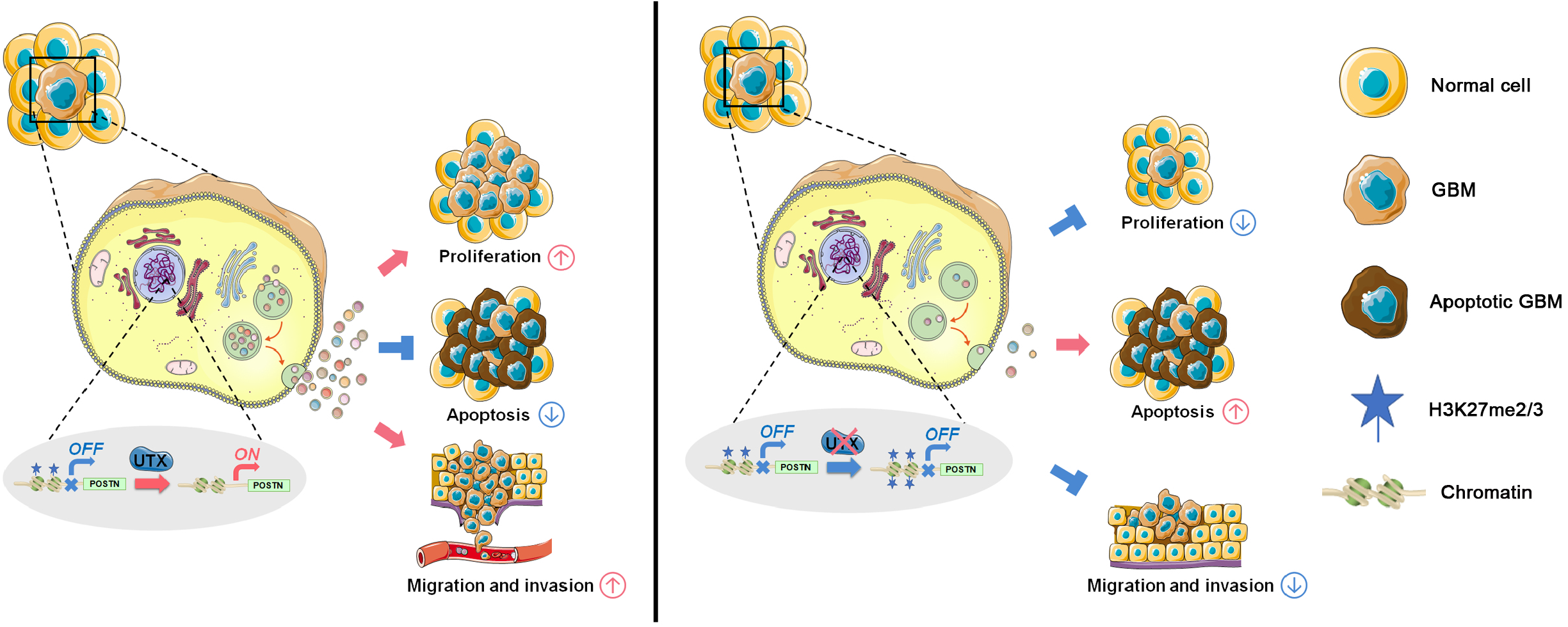
Schematic model of the mechanism by which UTX knockdown regulates glioblastoma progression. The expression level of UTX is significantly increased in GBM cells (left panel) and results in a low H3K27me2/3 level on the POSTN gene that culminates with activated transcription. POSTN protein could interact with lots of secreted proteins, including COL1A1, PTHLH, CPA4 and VCAM1. These proteins, the important components of the tumor microenvironment, are involved in the proliferation, apoptosis, migration and invasion in GBM cells. Knockdown UTX (right panel) significantly increases the H3K27me2/3 level of the POSTN gene, which in turn represses the transcription of POSTN. Fewer proteins, especially interaction with POSTN, are secreted to the extracellular matrix and change the tumor microenvironment, thereby inhibiting proliferation, migration and invasion, promoting cell apoptosis, even affecting proliferation and apoptosis of GSCs.

However, targeting UTX for cancer treatment remains controversial. Although lots of studies have provided evidence to show that UTX plays a pro-oncogenic role (Kaneko & Li, 2018), some studies have reported the role of UTX as a tumor suppressor (Ntziachristos *et al*, 2014; Xie *et al*, 2017). We showed that UTX is differentially expressed in a variety of tumors (Fig. EV5 A, B). This large difference may, in part, be responsible for the differences in the effect of UTX among different cancer types. Another reason may be that UTX is a subunit of MLL3 and MLL4, which are the members of the COMPASS family of histone H3 lysine 4 (H3K4) methyltransferases (Sze & Shilatifard, 2016). Compared to the UTX, H3K4 methylation has the opposite effect on the regulation of gene expression (Zaidi *et al*, 2013). H3K4 methylation is tightly associated with transcriptional start sites of actively transcribed genes (Shilatifard, 2012). The biological role of UTX and MLL3/MLL4 in cancer pathogenesis is quite complicated. The mechanistic relationship between UTX and MLL3/MLL4 and their regulation of enhancer activity, whether local or global is still unclear. Our research indicated that these chromatin proteins in the regulation of enhancer activity are not global in GBM, just few tumor suppressor genes were activated by UTX. Further studies are necessary to understand the fundamental mechanism and precise functional impact of MLL3/MLL4 and UTX alterations.

The TME is a dynamic network structure and has emerged as a key factor to regulate tumor progression (Ribeiro Franco *et al*, 2020; Wang *et al*, 2017). Therefore, anticancer research cannot be just understood the features of cancer cells, but instead should encompass the effect of the TME (Panigrahi *et al*, 2020). In this study, GO enrichment analysis showed that the genes of secretion, extracellular structure and matrix had changed after UTX knockdown. And a large number of ECM genes, including COL1A1, POSTN, VCAM1, PTHLH, and CPA4, actually changed which was detected by qRT-PCR, ELISA and Western blotting. These phenomena suggested that UTX mainly regulated GBM through alteration in the ECM. More importantly, protein-protein interactome analysis showed that POSTN protein could interact with most other proteins. As one of the matricellular proteins, POSTN is mainly secreted by stromal cells in normal tissues. But tumor cells, especially cancer stem cells, can also secret POSTN in solid tumors(Liu *et al*, 2019). POSTN has played a vital role in regulating tumor progression and tumorigenesis through remodeling various tumor microenvironments, such as cancer stem cell niche, perivascular niche, immunosuppressive microenvironment (Ghajar *et al*, 2013; Gonzalez-Gonzalez & Alonso, 2018; Masuoka *et al*, 2012; Zhou *et al*, 2015b). In this study, we demonstrated that replenishment of POSTN protein in the medium abolished the effect of UTX on proliferation and apoptosis in both GBM cells and patient-derived GSCs. These phenomena suggested that POSTN was a key regulator of the antitumor effect of UTX knockdown on GBM. However, the antitumor effect of UTX cannot simply account for the inhibition of POSTN expression. Many other tumor-associated proteins like EREG, ECSCR, STC2 and MPP4 were also regulated by UTX, we cannot rule out the effect of these proteins. Whether these proteins mediate the antitumor mechanism remains to be seen.

As the most malignant and prevalent primary brain tumor, GBM is a complex system and represents heterogeneity. Some with increased carcinogenicity, unlimited self-renewal potential, a higher capacity of stemness (Prager *et al*, 2019). These stem-like cells were considered as GSCs, one of the important origins of GBM relapse (Charles *et al*, 2011; Venere *et al*, 2011). Previous research demonstrated that GSCs displayed higher intrinsic chemo- and radioresistance (Lathia *et al*, 2015). Moreover, there is increasing evidence that chemoresistance and tumor recurrence are closely related to GSCs (Eramo *et al*, 2006). Therefore, a novel investigation of eradicating the GSCs is a therapeutic priority for GBM treatment. Our study showed that UTX knockdown inhibited proliferation while promoting the apoptosis of patient-derived GSCs. Remarkably, supplemental POSTN protein could partially abolish the effect of UTX on GSCs. It provided further evidence that UTX could be regarded as a novel and effective therapeutic target for GBM. Compared to the GSCs which sorting from cell lines, the patient-derived GSCs represent a more realistic response in terms of recapitulating the individual differences (Nagle *et al*, 2018). In this study, we used patient-derived GSCs to examine UTX functions, and this enhances the trustworthiness of our findings.

Tumor initiation and progression is a very complex process, genetic and epigenetic alterations are both associated with carcinogenesis (Thompson *et al*, 2018). The great potential for epigenetic regulation by histone methylation lies in the fact that histone methylation changes are reversible, allowing recovery of gene function with normal DNA sequences (Bennett & Licht, 2018). Our current study set out to investigate the impact of UTX on both GBM and GSCs progression and suggested that knockdown UTX exerted an anti-tumor effect on GBM. These results add to the rapidly expanding field of the effect of UTX and may contribute to the development of therapeutic strategies for GBM. However, additional research, especially using clinical samples, is necessary for an in-depth understanding of the precise mechanisms of UTX in GBM.

## Materials and Methods

### Cell culture

LN-229 and U251 MG cell lines were purchased from Procell (China) and identified by short tandem repeat (STR) analysis. 1×105 cells were seeded in T25 flasks and incubated in an incubator (SANYO, Japan) with 5% CO2 and 95% air at 37 °C. The medium consisted of Dulbecco’s Modified Eagle’s Medium (DMEM), 10% fetal bovine serum (FBS), 100 U/mL penicillin and 100 μg/mL streptomycin (all from Gibco, USA). The cell subculture was trypsinized, mechanically blown to create single-cell suspension, and cryopreserved with a liquid nitrogen tank when necessary.

Human glioblastoma stem-like cells (GSCs) were isolated from 3 post-surgical samples (GSC02, GSC05 and GSC08). GSC02 patient underwent surgery at the First Affiliated Hospital of Xi’an Jiaotong University; GSC05 patient underwent surgery at the Second Affiliated Hospital of Xi’an Jiaotong University; GSC08 patient underwent surgery at Shaanxi Provincial Peoples’s Hospital. None of the patients received any treatment prior to the intervention. This study was performed in accordance with the principles of the Declaration of Helsinki. Written informed consent was obtained before specimen collection. The detailed information was shown in Table EV1. Single GSCs were isolated as previously described and with minor modification (Azari et al, 2011). Briefly, after washing 3 times with cold DMEM/F12 (1:1) medium (Gibco), the GBM tissue was cut into small pieces and incubated with dissociation medium (200 μL Collagenase type I (10 mg / mL, Sigma), 200 μL Dispase II (20 mg / mL, Sigma) and 1.6 mL DMEM/F12 (1:1) medium) at 37 °C for 12 min. Then, the tissue was mechanically dissociated using a pipette and filtered using a 40 μm cell strainer (BD Falcon), followed by centrifugation at 1,000 rpm for 3 min. Cells were seeded at 200,000 cell/ml in non-adhesive T25 flasks with 5 mL complete medium, consisted of DMEM/F12 (1:1), 1% N2, 2% B27 (minus vitamin A), 20 ng/mL EGF, and 10 ng/mL bFGF. After culturing 5-7 days, the formation of 80-200 μM sphere was observed. For single-cell adhesive culture, the sphere was dissociated into single cells using ACCUTASE™ (Stemcell Technologies, Canada) and plated in poly-D-lysine-coated 24-well plates.

### Public data sets analysis

The TPM (transcripts per million reads) of UTX from TCGA cancers and matched TCGA normal and GETx data were visualized by GEPIA2 (http://gepia2.cancer-pku.cn/#analysis). For survival analyses, Hazard ratios (HR) and Kaplan–Meier plots were determined using GEPIA 2 (http://gepia2.cancer-pku.cn/#survival) based on the expression status of UTX in TCGA data sets.

### UTX knockdown treatment

The small interfering RNA (siRNA) specific to human UTX and scrambled siRNA negative control (siNC) were synthesized by Genechem (Shanghai, China). The sequences were as follows:

si-UTX-1: 5 ‘-AUUUCAGUGGGCUAUUAAATT-3’,
si-UTX-2: 5 ‘-ACGAAAUAUCAAGGUUUCATT-3’,
si-UTX-3: 5’-CUAUGGAUGCUUUGCAAGCTT-3’,
siNC: 5’-UUCUCCGAACGUGUCACGUTT-3’.

LN-229 and U251 MG cells were grown on 24-well plates, and Lipofectamine 2000 reagent (Invitrogen) was used to deliver the siRNA (100 nM). The knockdown efficiency was evaluated by RT-PCR and Western blot. The lentivirus vector containing shRNA targeting human UTX (shUTX) or negative control vectors (shNC) was purchased from Genechem (Shanghai, China). Two shNC lentivirus vectors were used in this study; one containing eGFP was used to measure transfection efficiency, and the other which did not express fluorescent protein was used for the majority of the subsequent experiment. (2 × 10^4^ cells/well) were seeded in a 24-well plate and infected with 2 μL shUTX (1 × 10^8^ virus, MOI = 1:10) or 1μl shNC (1 × 10^8^ virus, MOI = 1:5), respectively.

One day later, the infected cells were selected with puromycin at 24 hours. For the GBM cells, the concentration of puromycin was 2.5 μg/ml, and 1.6 μg/ml puromycin was used for GSCs.

### Immunostaining

Cells were plated onto the coverslips and fixed with 4% paraformaldehyde (PFA) at room temperature for 20 min followed by washing three times with PBS. Then, cells were permeabilized in 0.1% Triton X-100 for 15 min and blocked with blocking buffer (containing 5% bovine serum albumin and 5% horse serum in PBS) for 1 h, followed by incubating with the primary antibodies overnight at 4°C. After washing three times with PBS, cells were then incubated with suitable secondary antibodies. The information of the first and secondary antibodies were shown in Table EV2. The negative control samples were just incubated in the blocking buffer instead of the primary antibody. Nuclei were visualized with DAPI-containing mounting medium (Vector, USA). Images were acquired using a fluorescence microscope equipped with a digital camera (BX51 + DP71, Olympus, Japan) and analyzed with ImageJ software (NIH, USA). The sphere formation images were taken with a Leica SP8 confocal microscope equipped with a × 40 oil immersion lens.

### Flow cytometry analysis

For cell cycle analysis, the treated cells were dissociated with trypsin into single cells and fixed with pre-cooling 75% ethanol overnight at 4°C. After washing twice with PBS, cells were stained with Propidium Iodide solution (100 μg/mL, Sigma-Aldrich, USA) containing 100 μg/mL RNase A (New England Biolabs, USA) for 15 min at 37°C and away from light. The cell cycle analysis was performed using a FACSCalibur system (BD Biosciences, USA) with an excitation at 488 nm and emission at 630 nm. 1×105 cells were detected for each sample. The data were collected using the FACSortCellquect software (BD Biosciences), and the DNA content and cell cycle distribution were determined using the Modfit LT software (BD Biosciences). The proliferation index (PI) was used to evaluate the changes in the cell cycle distribution with the following formula: PI = (S+G2/M) / (G0/G1+S+G2/M).

Apoptosis analysis was performed using the FITC Annexin V apoptosis detection kit (BD Biosciences, USA). After treatment, cells were dissociated into single cells, washed twice with pre-cooling PBS and resuspended in binging buffer. Then, 200 μL of the cell suspension (more than 1×10^5^ cells) was transferred to a 5 mL FACS tube (BD Biosciences) and stained in duplicate with 10 μL of FITC Annexin V conjugate and 10 μL of propidium iodide (PI, 10 mg/mL) for 15 min in dark at room temperature. Apoptosis was analyzed using a FACSCalibur (BD Biosciences) with 4The data were collected with FACSort Cellquect software (BD Biosciences) and the percent of apoptotic cells was referred to the apoptotic index (AI) with the following formula: (LR+UR) / (UL+LL+LR+UR). Apoptosis was analyzed using a FACSCalibur (BD Biosciences) with 488 nm excitation for Annexin V (emission collected at 525 nm) and 488 nm excitation for PI (emission collected at 630 nm). The data were collected with FACSort Cellquect software (BD Biosciences) and the percent of apoptotic cells was referred to the apoptotic index with the following formula: (LR+UR) / (UL+LL+LR+UR).

### TUNEL assay

The terminal deoxynucleotidyl transferase dUTP nick end labeling (TUNEL) assay was used for detecting cell apoptosis according to the manufacturer’s instructions (Roche Diagnostics, USA). In brief, cells were fixed with 4% PFA for 30 min, followed by permeabilization using 0.1% Triton X-100 in 0.1% sodium citrate buffer for 2 min on ice. Then, cells were incubated with 50 μL TUNEL reaction mixture for 1 h at 37°C. After washing with PBS, nuclei were stained with DAPI-containing mounting medium (Vector). Images were acquired using a fluorescence microscope equipped with a 40× objective (BX51 + DP71, Olympus) and analyzed with an Image-Pro Plus 5.0 software (Media Cybernetics, USA).

### BrdU labeling

Following the treatment, GBM cells were incubated with 10 μg/mL of BrdU for 1 hour, and GSCs were treated with BrdU for 2 hours. The BrdU-labeled cells were further detected by immunostaining. To identify the BrdU-labeled cells, cells were pretreated with 2 N HCl for 30 min at 37°C, followed by neutralizing with 0.1 M borate buffer (pH8.5) for 15 min. The percentage of labeled cells was evaluated and normalized by the PI-stained nuclei or SOX2 positive cells.

### Western blot analysis

After the treatment, cells or tissues were collected and lysed in RIPA lysis buffer supplemented with Protease Inhibitor Cocktail (Roche, Germany) for 15 min on ice, followed by sonication (Sonics, USA) and centrifugation (Eppendorf, Germany). Then supernatants were collected, and protein concentration of the samples was measured using the BCA assay (Pierce, USA). After boiling with loading buffer, Proteins (20 μg – 40 μg depending on the target protein) were resolved by 10%-12% SDS-PAGE and transferred to polyvinylidene fluoride (PVDF) membranes (BioRad, USA). The membranes were blocked with 5% non-fat milk for 2 h at RT and subsequently probed with specific primary antibodies overnight at 4°C (Table EV2). After washing three times with TBST, the membranes were incubated with horseradish peroxidase-conjugated anti-rabbit or anti-mouse IgG for 2 h at room temperature. Then, immunoreactive bands were visualized with an enhanced chemiluminescent substrate according to the manufacturer’s protocol (Pierce). The bands were collected using GeneGnomeXRQ (Syngene, UK) and analyzed using the ImageJ 3.5 software. The expression levels of target proteins were determined and normalized to the housekeeping β-Actin. All Western blot data were presented in samples from at least 3 independent experiments.

### Colony formation assay

Cells (500 cells/well) were seeded into a six-well plate and cultured in the normal condition for 14 days. After fixation with 4% PFA, cells were stained with crystal violet solution. The number of colonies was counted for each sample and data were presented in samples from at least 3 independent experiments.

### Wound healing assay

Cells were seeded in a six-well plate and cultured until an 80-90% confluent monolayer was formed. The scratch a straight line in one direction scratches were made using 200μL pipette tips and the wells were washed twice with medium, cells were allowed to grow for an additional 24 h, photographs were taken on an inverted phase-contrast microscope (Olympus, Japan). The gap distance of the wound was measured by Image-Pro Plus 5.0 software, and the data were normalized to the average of the control.

### Transwell invasion analysis

The 8 μm transwell invasion chambers (Millipore) were pre-coated with 100 μL Matrigel (BD Biosciences). 2×10^4^ of Cells suspension were added in the upper Matrigel-coated chambers; 600 μL DMEM containing 10% FBS was filled with in the lower wells. After incubation for 24 h, cells were fixed and stained with 0.5 % violet crystal, and then cells on the upper portion of the membrane were removed with a cotton swab. The number of invaded cells was counted under a light microscope, and the data were normalized to the average of the control.

### Animal experiments

Pathogen-free male athymic BALB/c nude mice (5 weeks old) weighing 20-25 g were used for this study and all mouse experiments were approved by the Xi’an Jiaotong University Health Science Center Ethics Committee. The mice were purchased from the Xi’an Jiaotong University Laboratory Animal Center (Certificate No. 22-9601018). The staff at Xi’an Jiaotong University Laboratory Animal Center was responsible for housing and daily maintenance. Housing and environmental enrichment are according to standards. All efforts were undertaken to minimize the suffering of the mice. 100 μL of Normal LN229 (Blank), sh-NC-LN229 (shNC) or KD-UTX-LN229 (KD-UTX) suspension (1×10^7^ / mL) were implanted into the subcutis of the mice to establish the heterotopic xenograft model. Tumors were monitored every weekday and size was measured by using an electronic digital caliper. If the tumor size exceeds 2000 mm^3^ or the diameter exceeds 15 mm, the mice will be weeded out and euthanatized. Five mice were sacrificed at 3 d, 7 d, 14 d, and 28 d respectively. At each time point, tumors were isolated and tumor weight was measured, and tumor size was calculated by the formula: length×width^2^ ×0.52 (Tomayko & Reynolds, 1989). Subsequently, heterotopic tumor tissues were collected and used for Western bolt and ELISA analysis.

### RNA-sequencing (RNA-seq)

Total RNA was extracted using a Trizolreagent kit (Invitrogen) according to the manufacturer’s protocol. RNA quality was assessed on an Agilent 2100 Bioanalyzer (Agilent Technologies, Palo Alto, CA, USA) and checked using RNase-free agarose gel electrophoresis. Eukaryotic mRNA was enriched by Oligo(dT) beads, while prokaryotic mRNA was enriched by removing rRNA by Ribo-Zero™ Magnetic Kit (Epicentre, Madison, USA). Then the enriched mRNA was fragmented into short fragments using fragmentation buffer and reverse transcripted into cDNA with random primers. Second-strand cDNA was synthesized by DNA polymerase I, RNase H, dNTP and buffer. Then the cDNA fragments were purified with QiaQuick PCR extraction kit (Qiagen, The Netherlands), end-repaired, poly(A) added, and ligated to Illumina sequencing adapters. The ligation products were size selected by agarose gel electrophoresis, amplified by PCR and sequenced using Illumina HiSeq2500 by Gene Denovo Biotechnology Co. (Guangzhou, China). Raw data of RNA-seq reported in this study have been deposited in the Genome Sequence Archive (GSA) in BIG Data Center (https://ngdc.cncb.ac.cn/gsa-human/) under the accession number HRA001073. Bioinformatic analysis was performed using Omicsmart, a real-time interactive online platform for data analysis (http://www.omicsmart.com).

### Quantitative reverse-transcription PCR (qRT-PCR)

Total RNA was isolated from cells using Trizol reagent following the manufacturer’s instructions, and 2 μg RNA was reverse transcribed into cDNA using a RevertAid first-strand cDNA synthesis kit (ThermoFisher, USA) supplemented with Oligo(dT)18 and Random Hexamer Primer. qRT-PCR was performed with GoTaq^®^ qPCR Master Mix (Promega, USA) using an iQ5 Real-Time PCR Detection System (BioRad, USA). The primer pairs were synthesized by TaKaRa and displayed in Table EV3.

### Chromatin Immunoprecipitation (ChIP)-qPCR

ChIP was performed as previously described with minor modifications as detailed below (Lim *et al*, 2009). 100 μL single-cell suspensions (2×10^7^ / mL) were fixed by 1% formaldehyde for 10 min. Then, glycine solution (final concentration 125 mM) was added and incubated for 5 min. After twice washing with 500 μL ice-cold PBS, 300 μL lysis buffer were added and sonicated for 3 × 30s using VCX 500 (SONICS, USA). The lysate was diluted and incubated in Dynabeads™ Protein A (Invitrogen, USA) which had pre-incubated with 2 μg of H3K27me2/me3 antibody (39435, Active Motif, USA) or IgG (Abcam, UK). The immune complexes were incubated at 4°C, 20 rpm on a Tube Revolver Rotator for 12 hours. Then, the chromatin–antibody–bead complexes were washed 4 times in 100 μL ice-cold RIPA buffer, rinsed with 400 μL ChIP elution buffer (containing 50 μg RNase A) and incubated at 37 °C at 1,200 rpm on a Thermomixer (Eppendorf, Germany) for 1 h followed by incubating with 2 μL Proteinase K (New England Biolabs, USA) at 65 °C, 4 hours. An input control was processed in parallel. ChIP DNA was purified by phenol-chloroform isoamyl alcohol extraction, ethanol-precipitated and dissolved in 20 μL EB buffer (Qiagen, Germany). Analysis of DNA was performed using qRT-PCR with gene-specific primers Table EV4). Recovery of genomic DNA as the percentage input was calculated as the ratio of immunoprecipitate to input.

### The enzyme-linked immunosorbent assay (ELISA)

At the end of each time point, the cell culture medium was collected and analyzed using the POSTN, VCAM1 and Pro-Collagen I alpha 1 ELISA kit (R&D Systems) according to the manufacturer’s instructions. The absorbance was measured at 450 nm using a multimicroplate spectrophotometer (BioTek). Triplicate parallel wells were examined in all the experiments and the data were presented as the average of at least three independent experiments.

### Statistical Analysis

Statistical analyses were performed using the GraphPad Prism 5.0 software. Data were evaluated for normality and homogeneity of variance before comparison. Differences between groups were analyzed using one-way ANOVA, followed by Tukey’s post hoc test. The Kolmogorov-Smirnov test was used for normality and homogeneity. The data were shown as mean ± standard deviation, and *P* < 0.05 was considered as statistically significant difference.

## Author contributions

ZC. Zhang designed the experiments. XL. Chen, ZC. Zhang, and Y. Liu supervised the research. JW. Xue and K. Wang performed animal breeding. Y. Luan and YF. Liu performed most of the other experiments. Z. Zhang prepared manuscript drafts. KG. Ma and Y. Liu edited the paper.

## Acknowledgements

This work was supported by grants from National Natural Science Foundation of China (No. 81901156, 82001493); China Postdoctoral Science Foundation (2017M623152, 2018T111035, 2019M653662). The authors are thankful to Dr. Jia Wang, Dr. Zhenyu Guo and Dr Xingyu Miao for helping in collecting glioblastoma specimens.

